# Scalable composition frameworks for multicellular logic

**DOI:** 10.1101/150987

**Authors:** Sarah Guiziou, Pauline Mayonove, Federico Ulliana, Violaine Moreau, Michel Leclère, Jéróme Bonnet

**Affiliations:** Centre de Biochimie Structurale, INSERM U1054, CNRS UMR5048, University of Montpellier, France.; Laboratoire d’Informatique, de Robotique et de Microelectronique de Montpellier (LIRMM). CNRS UMR 5506, University of Montpellier, France.

## Abstract

A major goal of synthetic biology is to reprogram living organisms to solve pressing challenges in manufacturing, environmental remediation, or healthcare. While many types of genetic logic gates have been engineered, their scalability remains limited. Previous work demonstrated that Distributed Multicellular Computation (DMC) enables the implementation of complex logic within cellular consortia. However, current DMC systems require spatial separation of cellular subpopulation to be scalable to N-inputs, and need cell-cell communication channels to operate. Here we present scalable composition frameworks for the systematic design of multicellular consortia performing recombinase-based Boolean or history-dependent logic, and integrating an arbitrary number of inputs. The theoretical designs for both Boolean and history-dependent logic are based on reduced sets of computational modules implemented into specific cellular subpopulations which can then be combined in various manners to implement all logic functions. Our systems mark a departure from previous DMC architectures as they do not require either cell-cell communication nor spatial separation, greatly facilitating their implementation and making them fully autonomous. Due to their scalability and composability, we anticipate that the design strategies presented here will help researchers and engineers to reprogram cellular behavior for various applications in a streamlined manner. We provide an online tool for automated design of DNA architectures allowing the implementation of multicellular N-inputs logic functions at: http://synbio.cbs.cnrs.fr/calin.

## Introduction

Reprogramming the response of living cells to chemical or physical signals is a key goal of synthetic biology that would support the development of complex manufacturing processes, sophisticated diagnostics, or cellular therapies^1^. Researchers have engineered many types of Boolean logic gates operating in single cells by using transcriptional regulators^2–7^, RNA molecules^8, 9^, or site-specific recombinases^10–12^. Recombinases were also used to implement memory devices^13, 14^ and time-dependent logic systems that can track the order of occurrence of events or execute history-dependent gene expression programs^15–17^.

However, scaling-up single-cell logic systems requires solving multiple engineering challenges. First, when program complexity increases (number of inputs ≥ 3), the high number of parts needed can cause metabolic burden and affect cellular viability. Second, current design methods are mostly *ad-hoc*, and each logic function is implemented using a different genetic architecture that needs to be fully characterized and optimized to counter any context effects resulting from new arrangements of genetic components. Despite recent progresses towards predictable gate design^6^, some gates simply do not work or are too complex to be implemented within a single cell. In addition, in order to avoid cross-talk, single-cell logic systems need to use different components for every novel signal to be processed. While library of orthogonal regulatory components have greatly expanded in recent years^3, 5, 18, 19^, their deployment can be challenging and requires time-consuming optimization.

In nature, division of labor between cellular subpopulations is a ubiquitous mechanism allowing cellular communities to accomplish complex functions^20, 21^. Early efforts to engineer synthetic multicellular systems led to the construction of pulse generators^22^, pattern-forming communities^23^, predator-prey ecosystems^24^, synchronized oscillators^25, 26^, or distributed metabolic pathways^27^. Researchers also realized that problems faced by logic circuits operating in single cells could be addressed by distributing the logic program between different cells^28^. Because of the spatial separation allowed by cellular compartments, optimized regulatory components can be reused in different subpopulations. As the circuit is divided into smaller sub-circuits, metabolic burden is reduced. Finally, simple cellular computing modules can be composed in different manners and wired via cell-cell communication channels to obtain different logic functions. One example used spatially separated *E.coli* containing NOR gates and wired by quorum-sensing molecules to design all 2-input logic gates^29^. In this approach, the architecture strictly followed multilayered circuit designs inspired from electronics, in which only the cells in the last layer produce an output. However, engineering logic circuits within biological systems does not necessarily requires a strict transposition of electronic designs. Hence, specific features of biology could be used at our advantage to engineer logic systems in a more efficient manner^10, 28, 30^. Following this idea, researchers developed an alternative approach termed Distributed Multicellular Computing (DMC) to reliably implement complex genetic programs^28, 31–33^. DMC is based on the decomposition of a logic function into a sum of logic sub-functions, each performed by particular subpopulations of cells. DMC allows for the use of standard computational modules that can be combined in various manners to realize any given logic function of interest. Importantly, multiple cells are capable of producing the output which is therefore distributed among the cellular subpopulations. Recently, Macia and colleagues proposed a scalable composition framework to implement DMC within a multicellular consortia by using: (i) cellular computing units performing elementary IMPLY or NOT functions, (ii) cell-cell communication channels, and (iii) spatial separation^34^. While highly scalable and being able to execute complex logic programs using simple elements, this design requires spatial separation between each subpopulations, and therefore cannot function autonomously.

Here we present two composition frameworks enabling the systematic design of logic gates performing Boolean or history-dependent logic within an autonomous multicellular consortia. We designed our system using site-specific recombinases, more specifically serine integrases, which allow for flexible engineering of complex logic gates within single-layer genetic architectures^10, 11^. Because integrase recombination is irreversible in the absence of cofactors, our systems exhibit memory, are single use (“one-shot”) and belong to the family of asynchronous logic devices (i.e. the system can respond to multiple signals even if they do not arrive simultaneously).

Designs for both asynchronous Boolean logic and history-dependent logic are based on reduced libraries of cellular computing units responding to one or multiple inputs that can be composed at will to implement all desired logic functions (Fig. 1). We provide theoretical designs for N-inputs asynchronous Boolean logic gates, N-inputs events-order trackers, and up to 5 inputs gene-expression programs which behaviors depend on the order of occurrence of inputs.

**Figure 1.**
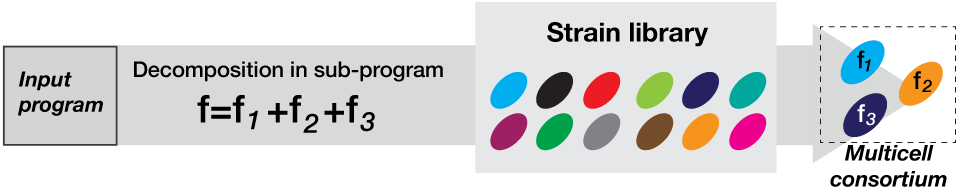
Distribution of logic programs within a multicellular consortia. The logic program of interest (either Boolean logic or history-dependent program) is decomposed as a disjunction (=sum) of sub-programs. A given function f encoding the program behavior is decomposed into functions *f*_1_, *f*_2_ and *f*_3_ encoding sub-programs. 3 strains performing *f*_1_, *f*_2_ and *f*_3_ are selected from the strain library to assemble a multicellular consortia computing the desired program.

Our systems mark a departure from previous DMC architectures as they do not require either cell-cell communication nor spatial separation, greatly facilitating their implementation and making them fully autonomous. We anticipate that these composable and scalable multicellular computing systems will support many applications requiring the implementation of complex genetic logic programs.

## Results

### Principle of multicellular asynchronous logic using integrase switches

In order to implement a logic program within a multicellular consortia, we decompose the program into several independent subprograms (or subfunctions)^34^. Each subprogram is then embedded within and executed by a different cellular subpopulation, chosen from a library containing a reduced number of cellular computing units (Fig. 1).

Several multicellular computing schemes have used cell-cell communication channels as chemical “wires” to connect the computations performed between different cellular sub-populations. However, such schemes can be less reliable in liquid culture^29^, and can limit future recomposition of the system. We thus designed our system so that each subcellular population computes independently of the others, with no cell-cell communication needed. As a consequence, in our system, if one of the subcellular population is ON (expression of the output gene), the global output of the system is considered to be ON. Because of their reduced number and of the absence of cell-cell communication, standard cellular computing units can be extensively characterized and optimized to predictably implement all desired logic programs within a multicellular consortia.

We designed our system to be operated using recombinases, more specifically serine integrases, which are members of the large serine recombinase family^35^ and perform site-specific recombination between attachment sites attB and attP. Recombination operates via a double-strand break located at the central dinucleotide followed by creation of hybrid sites attL and attR. Depending on the relative orientation of attB and attP, the recombination reaction leads to excision (parallel orientation) or inversion (antiparallel orientation) of the DNA sequence flanked by attachment sites^36^.

The robustness of serine integrases operation has enabled the engineering of many logic gates consisting of multiple recombination-sites disposed in various positions and orientations^10–12, 37^. The control signal driving integrase expression is decoupled from gate operation, so that gate architecture and components can be conserved even if the control signal is changed. Contrarily to multilayered gate architectures inspired from electronics, integrase-based logic supports the implementation of complex functions within a single genetic layer and with a reduced footprint^10, 11, 30^.

### A Hierarchical composition framework for asynchronous Boolean logic

Logic programs can be written as a Boolean equation (ex: *f*(*A*, *B*) = *A*.*B*). The Boolean equation corresponding to the output state (1 if output is on, 0 if off) is a function of the input states (1 if input has been present, 0 if input has never been present). We write Boolean functions using the disjunctive normal form (i.e. as a sum of products of input variables or their negations)^38^:

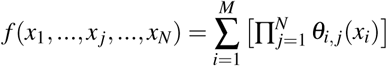

Following our program distribution in independent subprogram, each cellular computing unit performs a subprogram corresponding to a product of simple terms (part of the Boolean equation). The “sum” of products is performed by combining the various cellular computing units (Fig. 2A).

**Figure 2.**
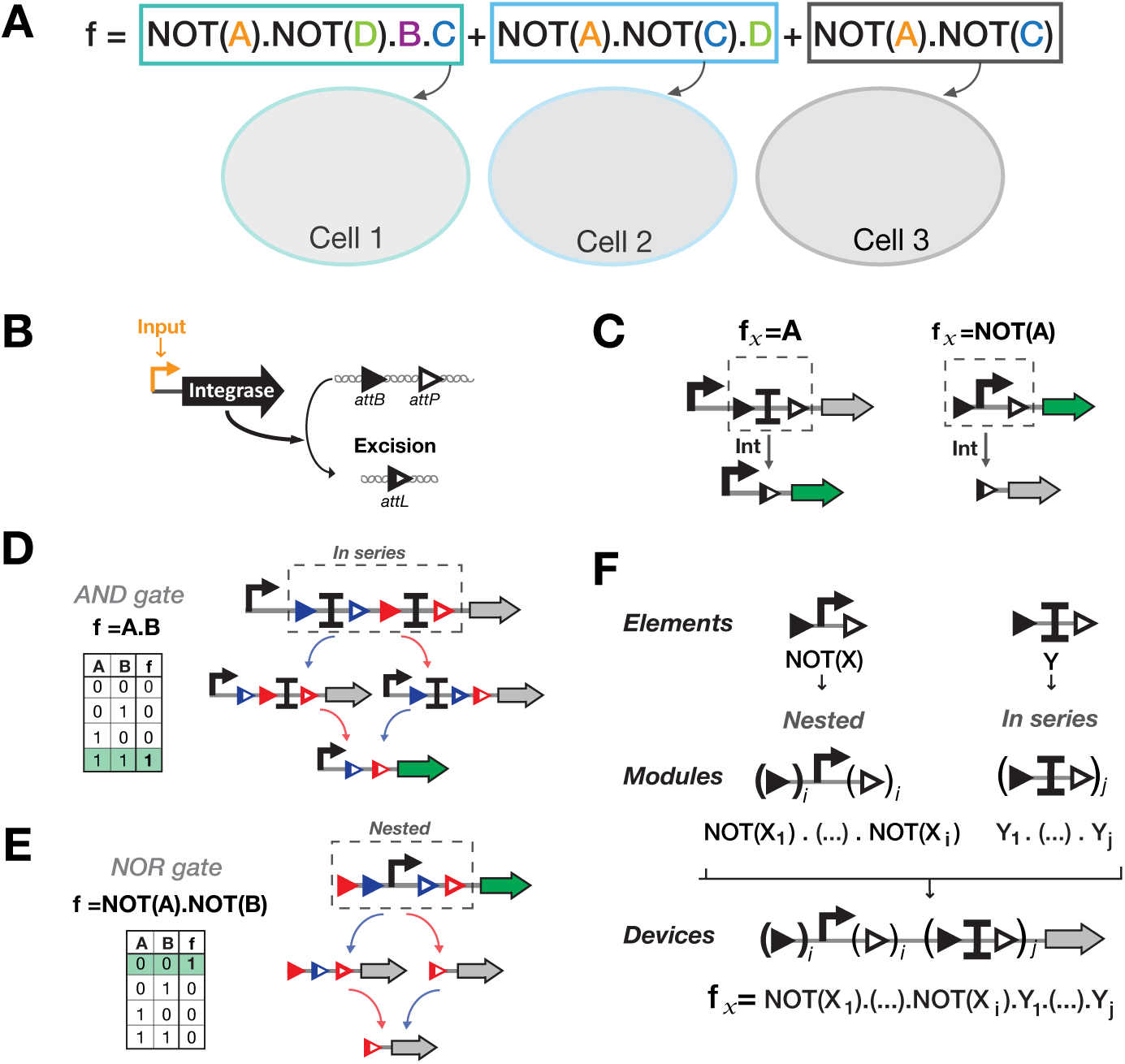
A Hierarchical composition framework for asynchronous Boolean integrase logic. **(A)** Distribution of a Boolean logic equation within a multicellular consortia. Boolean logic equations are decomposed into products of variables or their negations. Here an example is depicted in which a Boolean logic equation is decomposed into three sub-equations and implemented in three separate computing cellular units. **(B)** Principle of integrase-mediated excision. Integrase is expressed in response to a transcriptional signal and triggers excision of DNA sequences flanked by attB and attP disposed in parallel orientation. **(C)** Elementary computational elements. To obtain an IMPLY function, a transcriptional terminator is flanked by parallel att sites. In the absence of signal, transcription of the gene of interest is blocked. When the signal is present the terminator is excised, and the output gene is expressed. To obtain a NOT function, a promoter flanked by parallel att sites. In the absence of signal, the gene of interest is expressed. When the signal is present, the promoter is excised, and the gene is not expressed anymore. **(D)** Functional composition of IMPLY elements into IMPLY-s modules. IMPLY elements are composed in series to obtain product of IMPLY functions. For a 2-inputs IMPLY module, the output gene is expressed only when both inputs have been present and both terminators excised (corresponds to an AND gate (A.B)). **(E)** Functional composition of NOT elements into NOT modules. NOT elements are nested to obtain products of NOT functions. For a 2-inputs NOT module, the output gene is expressed only when none of the inputs has been present (corresponding to a NOR gate not(A).not(B)). **(F)** Hierarchical composition framework for Boolean integrase logic. IMPLY-s and NOT-s modules composed from elements are composed in series, following a priority rule in which the NOT-s module is placed upstream the IMPLY-s module. The device shown here can be scaled to all logic functions based on product of NOT and IMPLY functions.

For design simplification and robustness, we designed switches controlled only via integrase-mediated excision (Fig. 2B). Excision-based design reduces the distance between gate promoter and the gene of interest (see below, nested design for NOT functions). In addition, as no asymmetric terminator is needed, those systems might be easier to port into higher organisms^12^. We used two basic computational elements, the NOT and IMPLY functions, to implement sub-programs within individual cells. We then composed these elements in different manners to perform products of simple terms (Fig. 2C-F). We designed an IMPLY element by placing a transcriptional terminator flanked by recombination sites between the promoter and the output gene (Fig. 2C Left panel). Thus, in the absence of control signal, the output gene is not expressed. In the presence of signal, the terminator is excised, and the output gene is expressed (Output state =1). We designed a NOT element composed of a promoter surrounded by recombination sites in parallel orientation (excision mode) (Fig. 2C, right panel). In state 0, the promoter drives transcription of the output gene (Output state = 1). When the control signal is present, the integrase is expressed, the promoter excised, and the output gene will not not be expressed anymore (Output state = 0). We then defined a hierarchical composition framework to combine computational elements into higher-order computational modules. Each element belonging to similar or different classes follows specific functional compositions rules (Fig. 2D-E, and corresponding Fig. legends).

Finally, computational devices performing subprograms (i.e. products of NOT and IMPLY functions) are obtained by combining IMPLY and NOT modules in series. The priority rule specify that NOT modules are positioned upstream IMPLY modules when the output gene is in 3’ position (Fig. 2F).

Following this hierarchical composition framework, all sub-programs consisting of products of variables and variable negations are implementable within a cellular computing unit. All programs are then executed by a multicellular consortia containing different cellular computing units.

### Implementing all N-inputs Boolean functions from a reduced set of computational devices

We aimed to reduce the number of computational devices in order to simplify our system and allow for extensive characterization of all components. As the connection between inducible promoters (control signals) and integrases is interchangeable, we decided to implement only one computational device per set of symmetric functions. For example, for the symmetric functions not(A).B and B.not(A) only the computational device corresponding to the former function is implemented, while the latter is achieved by exchanging control signals A and B between the 2 integrases (Fig. S1). By performing this simplification, we could reduce the number of computational devices from 26 to 9 computational devices for all 3-inputs Boolean logic gates (256 functions) and from 80 to 14 for all 4-inputs Boolean logic gates (65 536 functions) (Fig. 3A and Fig. S1). For every additional control signal (from N-1 to N), only N+1 novel computational devices are needed while the number of Boolean functions increase drastically. For example, 7 additional devices are needed to transition from 5 to 6-inputs (27 devices in total), enabling the implementation of 10^10^ additional Boolean logic gates (for a total of 10^19^) (Fig. 3B).

**Figure 3.**
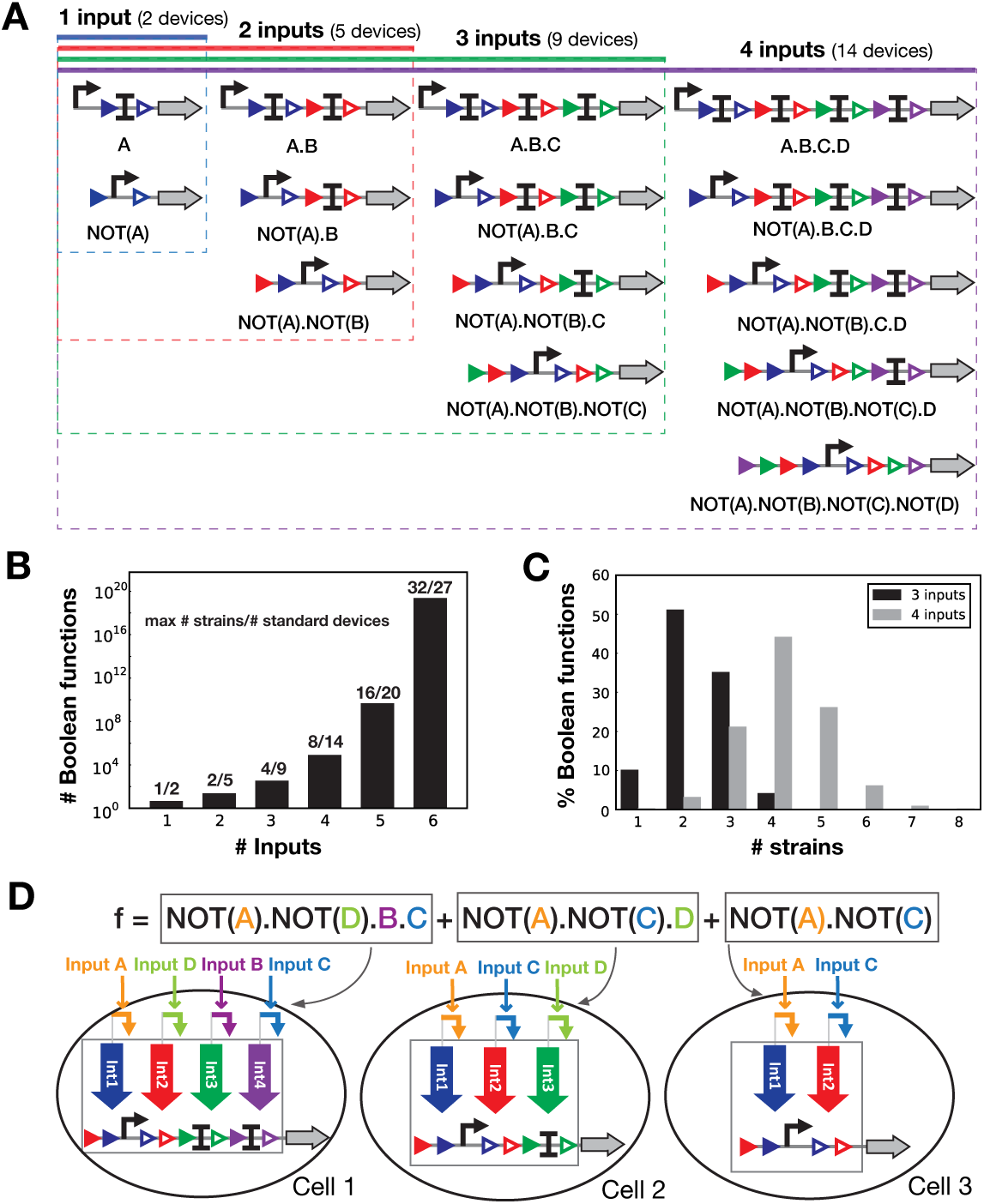
Implementing all Boolean logic functions using a reduced number of computational devices. **(A)** Schematics of all Boolean computational devices needed to implement for up to 4-inputs functions. We reduced the number of computational devices by connecting all signals to all integrases. Using this scheme, only one function per set of symmetric functions has to be encoded in a device. **(B)** Maximum number of strains and number of computational devices needed to compute all Boolean functions for a given number of inputs. See material and methods for details. **(C)** Proportion of Boolean functions implementable with a specific number of strains for 3 and 4-inputs. **(D)** Example of a biological implementation for a 4-inputs Boolean function. The function shown here is divided into sum of products (see Fig. 2A). Each product of simple terms is executed using previously defined computational devices into separated cellular computing units. By combining the different units, the full-logic function is obtained. If at least one of the cellular units is ON, the output is considered to be present. Of note, inputs are not always connected to the same integrase (as for input D in Cell1 and Cell2), and all integrases and inputs are not present in all Cells.

To realize N-inputs Boolean logic gates, a maximum of 2*^N^*^−1^ different cellular computing units have to be composed, corresponding to a culture of 2*^N^*^−1^ different strains: 4 for 3- inputs and 8 for 4-inputs (Fig. 3B). However, most gates can be composed using a reduced number of cellular computing units (an average of 2.3 strains for 3-inputs and 3.6 strains for 4-inputs gates) (Fig. 3C).

Importantly, using a multicellular system to perform Boolean logic programs reduces the size of genetic circuits embedded into individual strains. For a N-inputs Boolean equation, the different cellular computing units do not always comport N integrases and computational devices responding to N-inputs. As an example, the 4-inputs Boolean equation shown in Fig. 3D can be executed using 3 strains containing respectively 4, 3 and 2 integrases and with different control signals/integrases connectivity.

In summary, we designed a composable framework based on a reduced number of standard computational devices that support the systematic implementation of all N-inputs asynchronous Boolean logic gates within a multicellular population.

### Integrases-mediated memory for history-dependent multicellular behavior

By interlacing target sites for different recombinases, recombination reactions can be made dependent on each other. Using this concept, researchers started to implement genetic devices tracking the order of occurrence of signals, as well as history-dependent gene expression programs^15, 17, 39^, We found that a basic history-dependent motif could be repeatedly distributed into different cells to straightforwardly implement all input event-order trackers using a multicellular consortia (Fig. S2, Table S1). The state of the tracker could be addressed experimentally via multiplexed next-generation sequencing.

Previous methods designed history-dependent gene expression program using pairs of mutant recombination sites^17^. Finding a design to execute a given program relied on computational screening. Despite successful implementation of several 3-inputs programs, some programs were not accessible. In addition, the scalability of such systems might be challenging for several reasons. First, each program is executed using an ad-hoc design, requiring case-by-case optimization. Second, it is not clear how many pairs of mutant recombination sites can be used in parallel in a single-cell without any non-specific recombination reaction occurring. Third, repetitive DNA sequences often lead to genetic instability through homologous recombination^6, 40^ and fourth, highly-repetitive DNA sequences are notoriously difficult to synthesize.

We thus aimed at designing a composition framework enabling all possible history-dependent gene expression programs for up to 5 inputs to be systematically implemented within a multicellular consortia. To this aim, we used integrase switches performing site-specific DNA inversion and excision (Fig. 4A).

**Figure 4.**
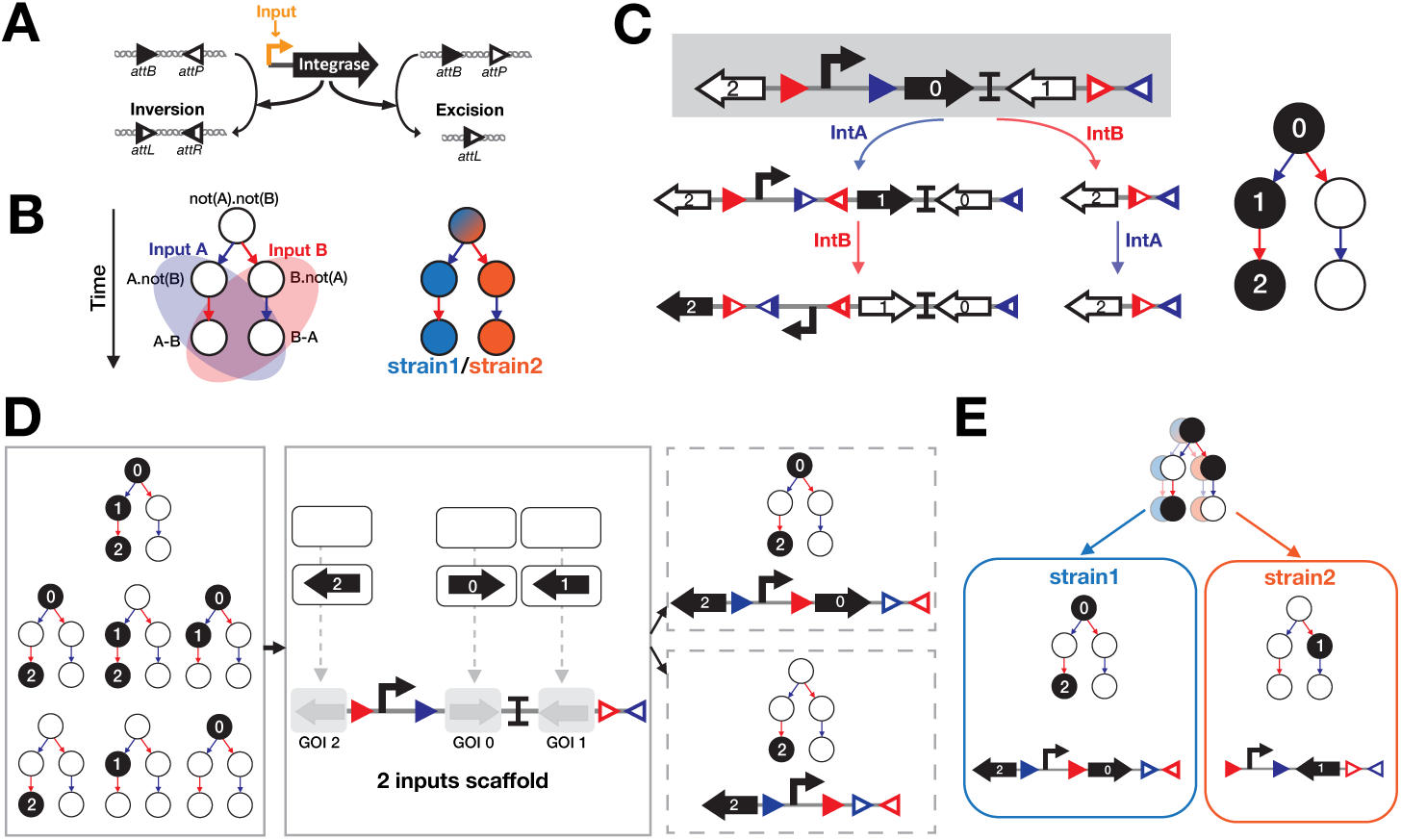
Modular scaffold to implement all 2-inputs history-dependent logic functions. **(A)** Integrase-mediated excision or inversion. When integrase sites are in the opposite orientation (left panel), the DNA sequence flanked by the sites is inversed. If integrase sites are in the same orientation (right panel), the DNA sequence flanked by the sites is excised. **(B)** Representation of history-dependent gene-expression programs and decomposition in separate strains. We represent history-dependent gene-expression programs as a lineage tree. Arrows represent the presence of inputs and nodes represent states of the system associated order-of-occurrence of the inputs. Gene-expression outputs associated to specific states are represented by a specific color or/and number. As the response of the system is history-dependent, the combinatorial state A.B (A and B) is separated in A then B (A-B) and B then A (B-A). For decomposition of the program into separate cells, each lineage of the history-dependent lineage tree is implemented in a different cell. **(C)** Operation of a 2-inputs history-dependent device. For each state of the lineage of interest, a different gene is expressed and no gene is expressed for states which are not in the lineage. Gene 0 is expressed when no input is present. If input A is present first, gene 1 is expressed, if input B which is present first, no gene is expressed (nor will be expressed) as the promoter is excised. If input B follows input A, gene 2 is expressed. **(D)** Gene swapping in a 2-inputs scaffold to obtain all possible programs for a single lineage. In the left panel, all 7 possible history-dependent programs for one lineage are listed. Output is equal to one in a state when the circle corresponding to this state is black. These programs are implementable based on our 2-inputs scaffold by addition of genes corresponding to the ON state in the corresponding GOI positions. In the right panel, two examples of biological implementation. For the first one, state 0 and 3 are ON, thus output genes are placed corresponding GOI positions. Same principle for the second example. **(E)** Example of implementation of 2-inputs history-dependent gene-expression programs in a multicellular consortia. As some states are ON in the two different lineages, each lineage is implemented using the 2-inputs scaffold with specific genes at appropriate positions.

### Modular scaffold designs for history-dependent gene expression programs

Each history-dependent gene expression program can be represented as a lineage tree (Fig. 4B for 2-inputs) in which each node corresponds to a state of the system (gene expression either ON or OFF) after inputs occurred in a particular sequence. Each lineage corresponds to a specific order-of-occurrence of the inputs. The number of lineages is equal to N! where N is the number of inputs. For instance, for 2-inputs, 2 lineages exist, while for 3 inputs, 6 lineages exist. In our design, we decompose the history-dependent gene expression program into subprograms corresponding to the different lineages. Each subprogram is then performed by a different cellular subpopulation (Fig. 4B right panel). Of note, we consider that the system operates in fundamental mode, i.e. inputs do not occur simultaneously, but sequentially.

We designed a modular scaffold supporting the execution of all 2-inputs history-dependent gene expression programs. The scaffold contains 3 directional cloning positions each supporting expression of a corresponding gene of interest (GOI) in a particular state of the lineage tree (Fig. 4C). Thus, any possible combination of gene expression states within a particular lineage can be achieved by simply inserting the desired gene at a given position (Fig. 4D). Importantly, depending on the identity of the different GOIs, the scaffold can be used to support single or multiple output programs.

Cellular subpopulations containing a scaffold incorporating different GOIs can be combined to perform multi-lineage history dependent genetic output (Fig. 4E). If control signals are exchanged between the different integrases, the same scaffold can be reused in all lineages. Following a similar principle, we designed scaffolds for 3, 4, and 5-inputs history dependent gene expression programs (Fig. 5A to E). The 3 and 4-inputs scaffolds allow for expression of a different GOI in each state of a given lineage (Fig. 5C for 3-inputs), while the 5-inputs scaffold allow expression of a different GOI in each states except the state 0 (no inputs). An additional strain is needed if gene expression is required in this condition.

**Figure 5.**
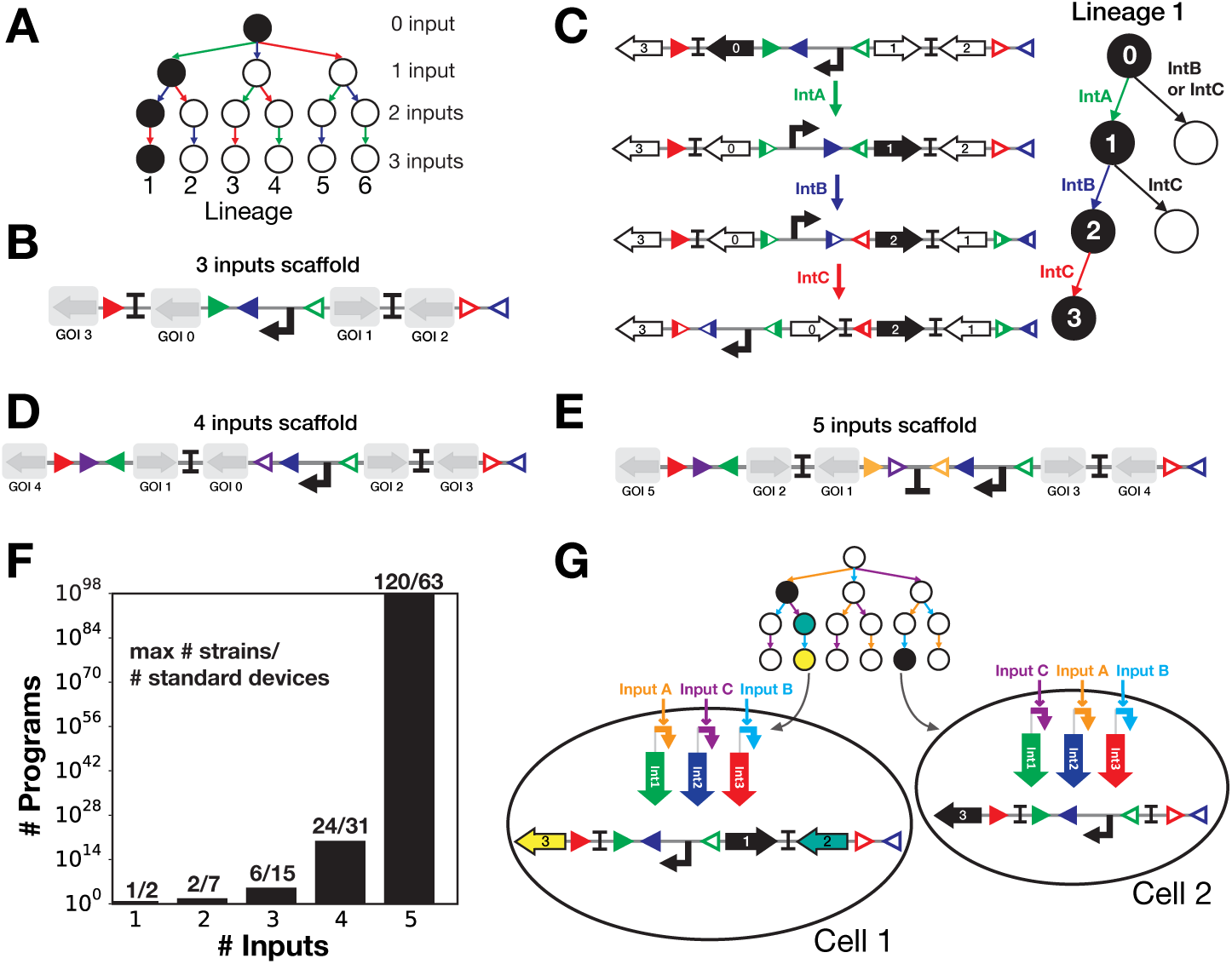
Scaling-up history-dependent scaffolds. **(A)** A 3-inputs lineage tree. The four rows of the lineage tree correspond to a different number of inputs that have occurred sequentially (from 0 to 3 inputs) and the 6 lineages to a different order-of-occurrence of inputs (example: A-B-C for lineage 1 and B-A-C for lineage 3). (B) 3-inputs scaffold design. Output genes are introduced at appropriate GOI positions (from 0 to 3) corresponding to the 4 states of a lineage (from 0 to 3 inputs occurrence) as in Fig 4. (C) Operation of the 3-inputs scaffold for one lineage. For each state of the lineage of interest, a different gene is expressed and no gene is expressed for states which are not in the lineage. Gene 0 is expressed when no input is present. **(D)** and **(E)** 4 and 5-inputs scaffold designs. Designs follow the same principles as in C. and Fig4. For the 5-inputs scaffold, no gene is expressed when no input is present. **(F)**. Maximum number of strains and number of computational devices needed to implement all programs for a given number of inputs and single output system (See material and methods for details). The graph represents number of possible history-dependent programs over the number of inputs. In order to reduce device number, all devices are based on a single scaffold and the different lineages are implemented by exchanging the signals driving integrase expression. **(G)** Example of 3-inputs and 3-outputs history-dependent gene-expression program. The lineage tree of interest shows 4 states ON with 3 different types of output in two different lineages. As two lineages exhibit an ON state, we implement this program in 2 different cell lines. The first cell computes the lineage A then C then B (Yellow-Purple-blue) with 3 ON states for different outputs. Consequently, different types of output genes are inserted in the corresponding GOI position. For the second cell, the lineage implemented is C then A then B (Purple-Yellow-blue), as an identical scaffold is used, inputs and integrases are connected to fit the lineage order.

The maximum number of cellular computation units needed to implement a history-dependent gene expression program is equal to the number of lineage (N! for N-inputs). For example, a maximum of 6 strains is needed for 3-inputs programs and 24 strains for 4-inputs programs (Fig. 5F). However, most functions are implementable with less than the maximum number of cells.

As for Boolean logic programs, a reduced number of standard devices is needed to implement all programs. Each N-inputs standard device corresponds to the N-inputs scaffold with insertion or not of genes at specific GOI position. To reduce the number of standard devices and to allow full-characterization of all devices, we chose to realize specific lineages by swapping connections between control signals and integrases instead of changing the position of integrase sites within the scaffold.

As an example for multiple-outputs programs, for a 3- inputs/3-outputs history-dependent gene-expression programs, 2 strains are needed, 3 different output genes are placed in the corresponding GOI positions and the 3-inputs are connected differently to integrases in the 2 different strains (Fig. 5G). If multiple outputs would need to be ON at the same time for a single state, the different output-genes could be positioned in a polycistronic architecture at the same GOI position. In summary, our scaffold-based design supports the execution of up to 5-inputs/N-outputs history-dependent gene-expression programs within a multicellular population.

## Discussion

In this work we developed scalable composition frameworks to implement asynchronous Boolean and history-dependent logic as well as N-inputs event-order trackers within a multicellular consortia. We provide an online (currently beta) design tool for the systematic design of asynchronous logic circuits called CALIN (Composable Asynchronous Logic using Integrase Networks). While these designs are currently theoretical, the robustness of integrase-mediated recombination against various sites permutations and orientations^10, 11, 17, 37^ should support straightforward experimental implementation.

As serine integrases do not require host-specific cofactors and can operate in many species, Boolean and history-dependent gene expression programs could be implemented in many organisms. For instance, history-dependent logic could be useful to control cellular differentiation^41^. Logic programs could also be distributed between different species operating in concert. In such schemes, researchers could take advantage of the particular capacities of different organisms to detect different signals and/or perform specific tasks. Example of applications include environmental remediation^42, 43^, or microbiome engineering for therapeutic applications^44^.

By taking advantage of the single-layer architecture of integrase logic, each subcellular population of the consortia performs complex logic functions. In consequence, the designs presented here exhibit two significant improvements over previous DMC systems: (i) no cell-cell communication channels (i.e. chemical “wires”) are needed, and (ii) cells do not need to be spatially separated, thereby supporting the implementation of fully autonomous multicellular consortia operating without external physical control device.

Another difference between our system and other DMC is the use of integrases switches that provide memory to the system and support time-dependent logic^15, 17, 39^. Because of recombinase mediated DNA data-storage, the state of the logic system can be not only be addressed through reporter gene expression but also via PCR or DNA sequencing^11, 15, 45^, even if the cells die. Such properties provide many flexible delayed readout modalities, and could be useful for applications like diagnostics or environmental monitoring.

As others DMC systems, for a given number of inputs, the number of elementary computational devices needed to compose all logic functions compares very favorably with the number of possible functions. For example, implementing all 65 536, 4 inputs, or all 4.10^9^, 5-inputs Boolean functions only requires 14 or 20 computational modules, respectively. For event-trackers, only 3 strains are sufficient to track the order of occurrence of 4 inputs (among 65 possible states, table S1), and 5 strains are required to track all 6-inputs sequence (among 1957 possible states). For history-dependent gene expression programs, 63 computational devices are sufficient to implement all 10^98^ possible 5-inputs history-dependent gene-expression programs, a number higher than the estimated number of atoms in the observable universe (10^79^,^46^) (Fig. 5F). Of note, some history-dependent logic programs (e.g. successive expression of a different gene at each stage of a specific input sequence) can be implemented by using only one scaffold in one cellular strain.

A possible limitation of our system is the high number of strains that has to operate in concert when the number of inputs increase (Fig. 3B). For example, some 4-inputs logic functions require up to 8 strains operating together. Co-cultivating such high-number of strains could cause several types of failures. For instance, cells expressing the output gene(s) could divide slower and be counter-selected from the population. This problem could be addressed through meticulous optimization of gene expression levels or by encapsulating the different strains into hydrogel beads^45^. Also, as the number of strains increase, the output of one subpopulation representing a small fraction of the whole consortia could become difficult to measure. Also, the output level in the global population will be different if one or multiple cellular subpopulation are turned ON. A cell-cell communication channel could be used to distribute the output within the whole-population (Fig. S3). Further work should thus be directed at finding efficient circuit minimization methods, enabling a reduction in the number of strains by exploiting existing redundancies within the different tree lineages (Fig. S4).

Finally, asynchronous Boolean logic might not be suited for applications requiring “real-time” response and reversibility. Interestingly, synchronous, real-time logic gates can be implemented based on reversible recombination reactions performed by integrases coupled with Recombination Directionality Factors (RDFs)^10, 30^. We thus also designed scalable hierarchical composition frameworks based on integrases/RDFs systems (Fig S5).

In conclusion, the scalable and composable designs presented here are a new addition to the toolbox of logic devices and will support research and engineering applications requiring complex programs to be executed by living cells.

## Acknowledgements

We thank L. Ciandrini, G. Cambray, J.L. Pons, G. Labesse, members of the synthetic biology group and of the CBS for fruitful discussions. Support was provided by an ERC Starting Grant, the INSERM Atip-Avenir program and the Bettencourt-Schueller Foundation. S.G. was supported by a PhD fellowship from the French Ministry of Research. The CBS acknowledges support from the French Infrastructure for Integrated Structural Biology (FRISBI) ANR-10-INSB-05- 01.

## Additional information

The CALIN webserver can be found at: http://synbio.cbs.cnrs.fr/calin. Supplementary materials are available online.

## Methods

### Equations for the determination of the number of functions/strains/devices for Boolean logic and History-dependent logic

For Boolean logic, the number of Boolean function corresponds to 2 to the power of the number of possible states, as each states can be equal to 1 or to 0. The number of possible states is equal to 2 to the power of N with N the number of inputs. Consequently, the number of Boolean logic function is equal to (eq1).

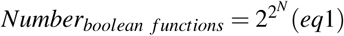

Then, the maximum number of strains needed to implement any Boolean logic function with N inputs is equal to (eq2), as all N-inputs Boolean equations can be written as a sum of 2*^N^*^−1^ product of variables or their negations^38^.

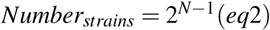

The number of different conjunctions (corresponding to a product of variables or their negations) is equal to (eq3).

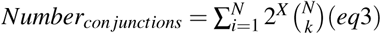

Then, if we implement all these functions within cells, the number of standard devices needed is equal to the number of conjunctions (eq4).

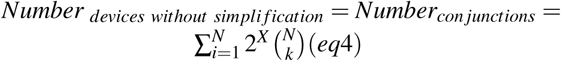

This method leads to a high number of devices. Therefore, we decided to construct only one device per set of symmetric functions (ex: *A*.*not*(*B*) is the symmetric function of *not*(*A*).*B*). This approach reduces the number of standard devices as in (eq5). In consequence, for a N-inputs function, devices computing from 1 to N-inputs are needed and N+1 non-symmetric function computing product of N-variables or its negation exist.

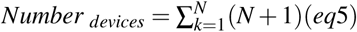

In first approximation, N sensor-modules in which a a control signal (i.e. a sensor device responding to an input of interest) is connected to an integrase are needed for the construction of a N-inputs system. However, as we reduced the number of devices to a set composed of non-symmetric functions, we need to connect all control signals to all integrases to enable all functions to be computed. Then, *N*^2^ sensor-modules are needed.

For history-dependent logic, the programs are represented as a lineage tree. Each node of this tree corresponds to a specific state of the system in response to different scenario: when none of the inputs occurred, when one input occurred, and when multiple inputs occurred in a particular sequence. For a N inputs-program, the number of states is equal to (eq6).

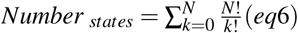

Then, for N-inputs 1-output history-dependent logic programs, the number of possible programs is equal to 2 to the power of the number of states (eq7), as all states can have either a ON or OFF output. Similarly for N-inputs M-outputs history-dependent logic programs, 2 to the power of the number of states multiplied by M programs exist (eq8).

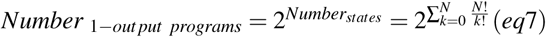

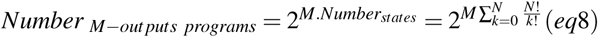

The maximum number of strains needed to implement a N-inputs/M-outputs history-dependent gene-expression program is equal to factorial of N which corresponds to the number of lineage in an N-inputs lineage tree.

The number of devices for 1-output/N-input system is equal to the number of states in one lineage and corresponds to the number of inputs present which can go from 0 input to N inputs (eq9).

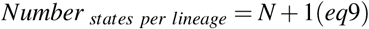

Consequently, the total number of devices needed to construct all N-inputs history-dependent programs is equal to (eq10).

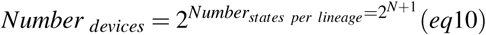

As for Boolean logic devices, we construct only standard history-dependent devices based on one scaffold and then differentially connect control signals to integrases to encode a specific lineage. The number of required sensor-modules is then identical than for Boolean logic.

### Automated generation of genetic designs to execute multicellular Boolean logic and History-dependent gene expression programs

We encoded algorithms for N-inputs Boolean logic circuit designs and up to 5-inputs History-dependent program designs using Python (Fig. S6). For both algorithms, the input of the program is a truth table, either a Boolean truth table or a lineage tree (equivalent to a sequential truth table). The output is the biological design, which corresponded to the number of strains needed, the design of each computing device and the specific integrases-inputs connections for each strain.

For the Boolean logic design (Fig. S6A), the truth table is transformed into a Boolean logic function in the disjunctive normal form using the Quine McCluskey algorithm^38^. We decompose the function in subfunction corresponding to the conjunctive terms (product of variables or variable negations). From each subfunction, we extract two types of informations. First, based on the numbers of IMPLY and NOT functions, we identify which Boolean logic devices are needed. Second, based on the association of inputs to either IMPLY and NOT functions, we identify which sensor-modules are needed among the different connection possibilities between control signals and integrases. Finally, by combining the designs determined for the different subfunctions, we obtain the global design for biological implementation of the desired truth table.

For History-dependent logic design (Fig. S6B), the lineage tree is decomposed in sub-tree composed of single lineage containing one or multiple ON states. This decomposition is done subtracting iteratively the lineages containing ON states. To obtain the lowest number of subprograms, we prioritize among the lineages with ON states the ones for which the highest number of inputs occured (from the right to the left of the lineage tree). After decomposition, for each selected lineage, two informations are extracted. First, based on which states are ON, we directly design the corresponding scaffold by specifically inserting genes at the adequate GOI positions. Second, the order-of-occurrence of inputs corresponding to the lineage is used to identify which sensor modules are needed among the different connection possibilities between control signals and integrases. Then, by combining the design of the different lineages, we obtain the global design for biological implementation of the desired history-dependent gene expression program.

As these methods are straightforward, they support the obtention, in a reduced time, of biological designs performing complex programs in response to a large number of inputs.

## References

1. Endy, D. Foundations for engineering biology. Nat. 438, 449–453 (2005).

2. Fischer, C. & Fussenegger, M. BioLogic gates enable logical transcription control in mammalian cells - kramer - 2004 - biotechnology and bioengineering - wiley online library. BioTechnology (2004).

3. Stanton, B. C. et al. Genomic mining of prokaryotic repressors for orthogonal logic gates. Nat. Chem. Biol. 10, 99–105 (2014).

4. Anderson, J. C., Voigt, C. A. & Arkin, A. P. Environmental signal integration by a modular AND gate. Mol. Syst. Biol. 3 (2007).

5. Moon, T. S., Lou, C., Tamsir, A., Stanton, B. C. & Voigt, C. A. Genetic programs constructed from layered logic gates in single cells. nature.com 491, 249–253 (2012).

6. Nielsen, A. A. K. et al. Genetic circuit design automation. Sci. 352, aac7341 (2016).

7. Ausländer, S., Ausländer, D., Müller, M., Wieland, M. & Fussenegger, M. Programmable single-cell mammalian biocomputers. Nat. 487, 123–127 (2012).

8. Win, M. N. & Smolke, C. D. Higher-Order cellular information processing with synthetic RNA devices. Sci. 322,456–460(2008).

9. Lucks, J. B., Qi, L., Mutalik, V. K., Wang, D. & Arkin, A. P. Versatile RNA-sensing transcriptional regulators for engineering genetic networks. Proc. Natl. Acad. Sci. U. S. A. 108, 8617–8622 (2011).

10. Bonnet, J., Yin, P., Ortiz, M. E., Subsoontorn, P. & Endy, D. Amplifying genetic logic gates. Sci. 340, 599–603 (2013).

11. Siuti, P., Yazbek, J. & Lu, T. K. Synthetic circuits integrating logic and memory in living cells. Nat. Biotechnol. 31, 448–452 (2013).

12. Weinberg, B. H. et al. Large-scale design of robust genetic circuits with multiple inputs and outputs for mammalian cells. Nat. Biotechnol. 35, 453–462 (2017).

13. Podhajska, A. J., Hasan, N. & Szybalski, W. Control of cloned gene expression by promoter inversion in vivo: construction of the heat-pulse-activated att-nutL-p-att-N module. Gene 40, 163–168 (1985).

14. Bonnet, J., Subsoontorn, P. & Endy, D. Rewritable digital data storage in live cells via engineered control of recombination directionality. Proc. Natl. Acad. Sci. U. S. A. 109, 8884–8889 (2012).

15. Ham, T. S., Lee, S. K., Keasling, J. D. & Arkin, A. P. Design and construction of a double inversion recombination switch for heritable sequential genetic memory. PLoS One 3, e2815 (2008).

16. Friedland, A. E. et al. Synthetic gene networks that count. Sci. 324, 1199–1202 (2009).

17. Roquet, N., Soleimany, A. P., Ferris, A. C., Aaronson, S. & Lu, T. K. Synthetic recombinase-based state machines in living cells. Sci. 353, aad8559 (2016).

18. Rhodius, V. A. et al. Design of orthogonal genetic switches based on a crosstalk map of σs, anti-σs, and promoters. Mol. Syst. Biol. 9,702 (2013).

19. Yang, L. et al. Permanent genetic memory with ¿1-byte capacity. Nat. Methods 11, 1261–1266 (2014).

20. Hays, S. G., Patrick, W. G., Ziesack, M., Oxman, N. & Silver, P. A. Better together: engineering and application of microbial symbioses. Curr. Opin. Biotechnol. 36, 40–49 (2015).

21. Wintermute, E. H. & Silver, P. A. Dynamics in the mixed microbial concourse. Genes Dev. 24, 2603–2614 (2010).

22. Basu, S., Mehreja, R., Thiberge, S., Chen, M.-T. & Weiss, R. Spatiotemporal control of gene expression with pulse-generating networks. Proc. Natl. Acad. Sci. U. S. A. 101, 6355–6360 (2004).

23. Basu, S., Gerchman, Y., Collins, C. H., Arnold, F. H. & Weiss, R. A synthetic multicellular system for programmed pattern formation. Nat. 434, 1130–1134 (2005).

24. Balagaddé, F. K. et al. A synthetic escherichia coli predator-prey ecosystem. Mol. Syst. Biol. 4, 187 (2008).

25. Danino, T., Mondragón-Palomino, O., Tsimring, L. & Hasty, J. A synchronized quorum of genetic clocks. Nat. 463, 326–330 (2010).

26. Prindle, A. et al. A sensing array of radically coupled genetic’biopixels’. Nat. 481, 39–44 (2011).

27. Shong, J., Jimenez Diaz, M. R. & Collins, C. H. Towards synthetic microbial consortia for bioprocessing. Curr. Opin. Biotechnol. 23, 798–802 (2012).

28. Macía, J., Posas, F. & Solé, R. V. Distributed computation: the new wave of synthetic biology devices. Trends Biotechnol. 30, 342–349 (2012).

29. Tamsir, A., Tabor, J. J. & Voigt, C. a. Robust multicellular computing using genetically encoded NOR gates and chemical ‘wires’. Nat. 469, 212–215 (2011).

30. Subsoontorn, P. Reliable Functional Composition of a Recombinase Device Family, Https://purlStanfordEdu/sm186rb. Ph.D. thesis (2014).

31. Regot, S. et al. Distributed biological computation with multicellular engineered networks. Nat. 469, 207–211 (2010).

32. Macia, J. & Sole, R. How to make a synthetic multicellular computer. PLoS One 9, e81248 (2014).

33. Goñi-Moreno, A., Amos, M. & de la Cruz, F. Multicellular computing using conjugation for wiring. PLoS One 8, e65986 (2013).

34. Macia, J. et al. Implementation of complex biological logic circuits using spatially distributed multicellular consortia. PLoS Comput. Biol. 12, e1004685 (2016).

35. Groth, A. C. & Calos, M. P. Phage integrases: biology and applications. J. Mol. Biol. 335, 667–678 (2004).

36. Grindley, N. D. F., Whiteson, K. L. & Rice, P. A. Mechanisms of site-specific recombination. Annu. Rev. Biochem. 75,567–605 (2006).

37. Rubens, J. R., Selvaggio, G. & Lu, T. K. Synthetic mixed-signal computation in living cells. Nat. Commun. 7,11658 (2016).

38. Enderton, H. & Enderton, H. B. A Mathematical Introduction to Logic (Academic Press, 2001).

39. Hsiao, V., Hori, Y., Rothemund, P. W. & Murray, R. M. A population-based temporal logic gate for timing and recording chemical events. Mol. Syst. Biol. 12, 869 (2016).

40. Sleight, S. C. & Sauro, H. M. Visualization of evolutionary stability dynamics and competitive fitness of escherichia coli engineered with randomized multigene circuits. ACS Synth. Biol. 2, 519–528 (2013).

41. Brambrink, T. et al. Sequential expression of pluripotency markers during direct reprogramming of mouse somatic cells. Cell Stem Cell 2, 151–159 (2008).

42. Li, L. et al. Removal of methyl parathion from artificial off-gas using a bioreactor containing a constructed microbial consortium. Environ. Sci. Technol. 42, 2136–2141 (2008).

43. De Lorenzo, V. Systems biology approaches to bioremediation. Curr. Opin. Biotechnol. 19, 579–589 (2008).

44. Mimee, M., Citorik, R. J. & Lu, T. K. Microbiome therapeutics - advances and challenges. Adv. Drug Deliv. Rev. 105,44–54 (2016).

45. Courbet, A., Endy, D., Renard, E., Molina, F. & Bonnet, J. Detection of pathological biomarkers in human clinical samples via amplifying genetic switches and logic gates. Sci. Transl. Med. 7 (2015).

46. Eddington, A. S. The constants of nature. In Schuster, S. (ed.) The world of mathematics 2, vol. 2, 1074–1093 (1956).

